# Angiotensin receptor conformations stabilized by biased ligands differentially modulate β-arrestin interactions

**DOI:** 10.1101/2025.05.28.656666

**Authors:** Matthias Elgeti, Julia Belyaeva, Mahdi Bagherpoor Helabad, Dean P. Staus, Laura M. Wingler

## Abstract

“Biased” ligands of the angiotensin II type 1 receptor (AT1R) preferentially activate G protein or β-arrestin pathways by stabilizing distinct receptor conformations. Here we show that β-arrestin-biased AT1R ligands further vary in their ability to stabilize different modes of β-arrestin interaction, specifically interactions with the AT1R seven-transmembrane core versus the phosphorylated C-terminus. By combining biochemical assays with double electron-electron resonance (DEER) spectroscopy and integrative modeling, we show that ligands less effective at stabilizing the core complex promote an AT1R conformation with an intermediate transmembrane helix 6 position that is incompatible with β-arrestin core binding. Since the core and phosphosite interactions differentially activate the signaling, internalization, and desensitization functions of β-arrestin, our data demonstrate that the allosteric effects of GPCR ligands could directly modulate β-arrestin activities. This “intra-transducer bias,” or bias toward various functions of the same transducer, could enable finer control of GPCR drugs’ pharmacology than previously thought possible.

## Introduction

β-Arrestins are key transducers of G protein-coupled receptors (GPCRs), which they engage through multiple binding modes^1,2^. β-Arrestins can interact with GPCRs through phosphorylated regions in the GPCR C-terminus or intracellular loop 3 (ICL3) (“phosphosite interactions”) and through the GPCR seven-transmembrane (7TM) core, the same region to which heterotrimeric G proteins bind. Besides these two principal interfaces, β-arrestins have also been observed to interact with the cytosolic Helix 8 (H8) of the glucagon receptor^3^ and with membrane lipids^3–6^. GPCR/β-arrestin complexes can equilibrate between conformations where β-arrestin is engaged with only the phosphosites or with both the phosphosites and the 7TM core^7^. This equilibrium has important implications because each interaction is associated with different functions of β-arrestins. While the phosphosite interactions are sufficient to promote GPCR internalization and initiate many axes of β-arrestin-dependent signaling, the 7TM core interaction is required for certain signaling functions and the steric blockade of G protein activation^8–10^.

“Biased” GPCR ligands preferentially activate G proteins, GPCR kinases (GRKs), or β-arrestins^11^. The discovery of biased signaling has galvanized efforts to develop next-generation GPCR drugs that stimulate desired therapeutic effects but not on-target side effects. For example, upon binding to the extracellular ligand-binding site (“orthosteric site”) of the angiotensin II type 1 receptor (AT1R), angiotensin II (AngII) activates both vasoconstrictive Gq signaling and cardioprotective β-arrestin pathways^12–15^. β-arrestin-biased AngII analogs, which activate β-arrestin-mediated but not Gq-mediated responses, have been investigated as alternatives to clinically used AT1R antagonists for acute heart failure^16,17^ and acute respiratory distress syndrome associated with COVID-19^18^, and they continue to be of interest for indications such as dilated cardiomyopathy^19^ and aortic aneurysm^20^. While most studies of biased ligands have focused on differential activation of G proteins versus β-arrestins, more nuanced pharmacology has also been observed, including ligand biases toward G protein subtypes^21^ and activation of GRKs without subsequent receptor phosphorylation and β-arrestin recruitment^22^.

Since the physiological functions of β-arrestin are separable and encoded by different complex conformations, it is possible that ligands could further display “intra-transducer bias,” or selective modulation of various β-arrestin activities. Assessing ligand effects on the conformations of GPCR/β-arrestin complexes is challenging because most conventional readouts of β-arrestin activation reflect the phosphosite interactions. Accordingly, we developed an *in vitro* method to form AT1R/β-arrestin complexes mediated only by the 7TM core. We find that β-arrestin-biased ligands vary dramatically in their ability to stabilize such complexes. Double electron-electron resonance (DEER) distance measurements and MD simulation data provide a mechanistic explanation for why β-arrestin cannot robustly interact with one of the intracellular conformations stabilized by β-arrestin-biased ligands. These findings demonstrate that orthosteric ligands can allosterically regulate the conformational equilibrium of GPCR/β-arrestin complexes, which opens additional dimensions for refining GPCR drug pharmacology.

## Results

### β-arrestin1 interacts with the AT1R 7TM core in a ligand-dependent manner

We developed a strategy to investigate the interaction of β-arrestin with the AT1R 7TM core. To activate β-arrestin, we purified a complex of β-arrestin1, a synthetic phosphopeptide from the C-terminus of the V2 vasopressin receptor (V2R), and conformationally selective antibody fragments^4,23,24^ **(Fig. 1A, Fig. S1)**. These binding partners effectively promote conformational changes in β-arrestin1 that facilitate its interaction with the 7TM core of GPCRs, including interdomain rotation and stabilization of the extended finger loop^23^.

**Fig. 1.**
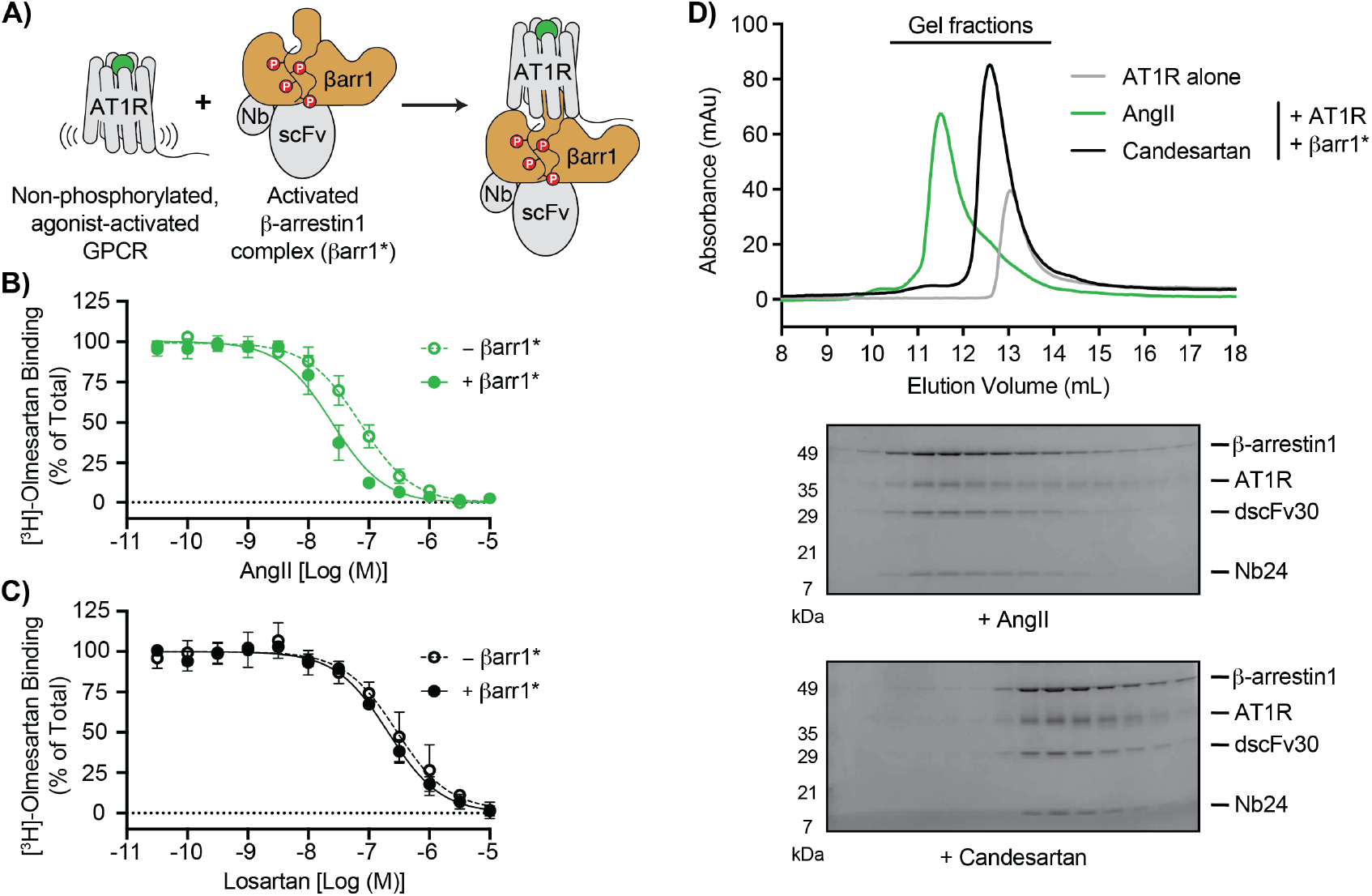
Formation of 7TM core-mediated AT1R/β-arrestin complexes. **(A)** Schematic of complex formation strategy. Purified, non-phosphorylated AT1R bound to various orthosteric ligands is incubated with βarr1*, a purified complex of β-arrestin1, the synthetic V2R phosphopeptide (V2Rpp), and the active-state-specific β-arrestin1 binders dscFv30 (or Fab30) and Nb24. **(B**,**C)** Competition binding between the radiolabeled antagonist [^3^H]-olmesartan and unlabeled (B) AngII or (C) losartan at purified AT1R in the absence or presence of βarr1* (2.5 μM). Log IC50 values are provided in **Table S1**. Data represent the mean ± S.D. of 3 independent experiments. **(D)** SEC and gel electrophoresis analysis (Coomassie staining) of deglycosylated AT1R/βarr1* complexes formed in the presence of AngII or candesartan.

Agonist-induced conformational changes in GPCRs increase their affinity for transducer proteins, and transducers reciprocally increase the affinity of GPCRs for agonists. Accordingly, we performed competition radioligand binding to determine whether the activated β-arrestin1 complex (βarr1*) recognizes AngII-stabilized AT1R conformations. βArr1* increases the affinity of AngII for purified, non-phosphorylated AT1R in an MNG micellar environment, but it does not alter the binding of the antagonist losartan **(Fig. 1B,C, Table S1)**.

We then used size exclusion chromatography (SEC) to evaluate whether ligands could promote stable complex formation between AT1R and βarr1*. AngII-bound AT1R co-migrates with βarr1*, as evidenced by a reduced retention time **(Fig. 1D)**. In contrast, complexes formed with the high-affinity antagonist candesartan dissociate. These data demonstrate that AT1R and βarr1* productively interact in an agonist-dependent manner through the receptor’s 7TM core.

### Intracellular conformation of βarr1*-bound AT1R

We previously used double electron-electron resonance (DEER) to obtain a detailed picture of how AngII, antagonists, and biased ligands alter the conformational heterogeneity of the intracellular regions of AT1R^25^. This method allows us to measure the distribution of distances between ten spin label pairs that cover the cytoplasmic receptor surface, including sites in TM1, TM5, TM6, TM7, ICL2, and H8 **(Fig. 2A)**. To understand how βarr1* affects the intracellular regions of AT1R, we acquired DEER data for each of these pairs in the presence of excess AngII and βarr1*.

**Fig. 2.**
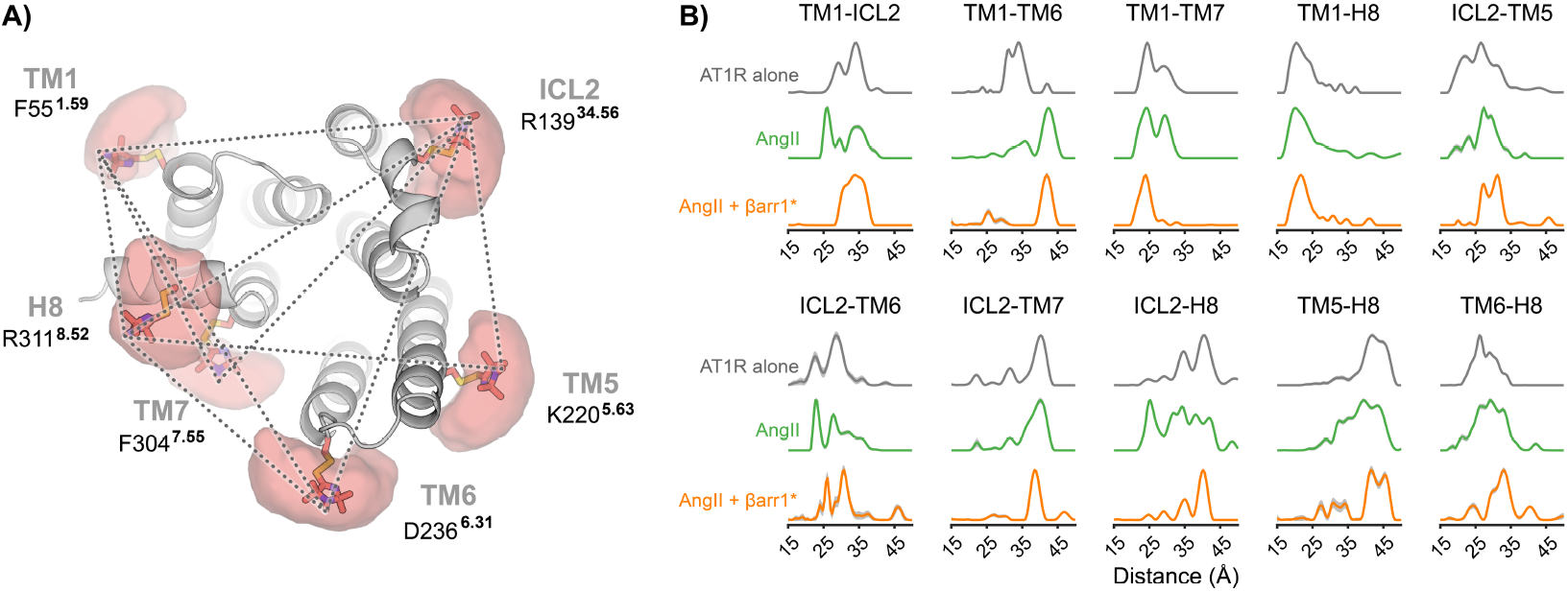
DEER analysis of AT1R activation and βarr1* core interaction. (**A**) Spin label positions and pairwise combinations (dashed lines) used to record distances in this study. Spin labels are shown as sticks; clouds indicate the possible positions of the unpaired electron. **(B)** Probability distance distributions between the indicated spin pairs showing the effects of AngII (green), and AngII + βarr1* (orange) on apo AT1R (gray). Shaded areas indicate 95% confidence intervals.

As previously reported^25^, AngII populates a heterogeneous ensemble of conformations, some of which exhibit hallmarks of the canonical active conformation of GPCRs **(Fig. 2B)**. These include the outward displacement of TM6 (cf. TM1-TM6 distances) and the inward movement of TM7 (cf. ICL2-TM7 distances). Addition of βarr1* markedly reduces this conformational heterogeneity **(Fig. 2B)**. The TM1-TM6 distribution shows that βarr1* stabilizes an active conformational state of TM6 (~42 Å). A smaller peak at ~25 Å also appears, which we assign to an active conformation with a rotated position of TM6, as previously observed for rhodopsin-transducin complexes^26^. Most pairs involving the ICL2 and H8 labeling sites show two major populations in the presence of βarr1*, indicating that flexibility remains in these regions.

### β-arrestin-biased ligands differentially stabilize AT1R/βarr1* complexes

TRV023, TRV026, and TRV027 are well-characterized biased ligands that recruit β-arrestin to the plasma membrane with almost indistinguishable potencies and efficacies **(Fig. 3A)**, consistent with previously reported data^27,28^. Even though their pharmacological profiles are similar, our previous DEER data showed these β-arrestin-biased ligands stabilize intracellular conformational distributions that are not only different from other ligand classes but also different from each other^25^ **(Fig. 3B, Fig. S2)**. In the main conformation stabilized by TRV026, designated “Occluded 1,” TM6 moves to an intermediate (TM1-TM6 35 Å) rather than the fully outward position (TM1-TM6 42 Å), but the conformation does exhibit a characteristic inward movement of TM7 (cf. ICL2-TM7 distances) and a substantial movement of H8 (cf. ICL2-H8 and TM5-H8 distances). In the main conformation stabilized by TRV023, designated “Occluded 2,” TM6 moves to the fully outward position (TM1-TM6 42 Å). However, mapping of the characteristic peaks indicates that TM5 does not rotate to the same extent as the fully activated conformation stabilized by AngII, and ICL2 undergoes a major conformational change^25^. TRV027 stabilizes approximately equal populations of Occluded 1 and Occluded 2^25^.

**Fig. 3.**
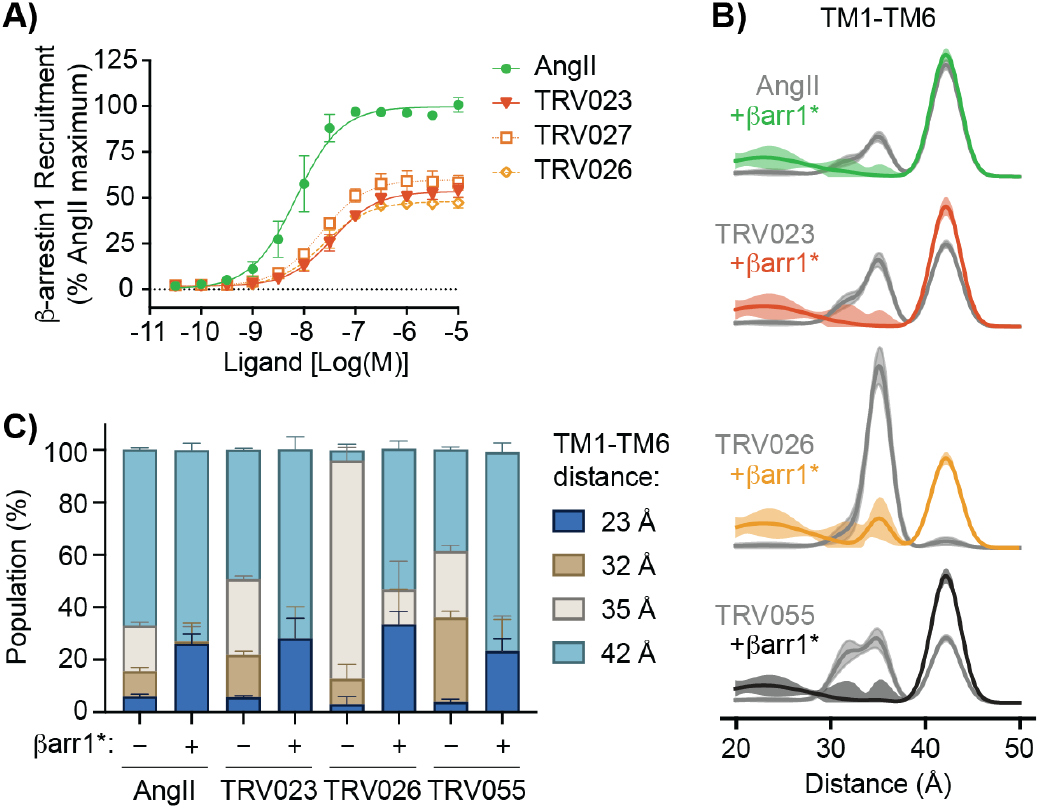
Formation of AT1R/βarr1* complexes by β-arrestin-biased ligands. **(A)** Ligand-induced β-arrestin1 recruitment to the cell plasma membrane. Data are normalized to the AngII maximum response and represent the mean ± S.D. of 3 independent experiments. Curve analysis parameters are provided in **Table S2. (B)** TM1-TM6 probability distance distributions for indicated ligands in the absence (gray) and presence (colored) of βarr1* using a Gaussian mixture model. Shaded areas indicate 95% confidence intervals. The associated raw DEER data and model-based analysis are shown in **Fig. S3. (C)** Median populations ± 95% confidence intervals of Gaussian populations centered around 23, 32, 35, and 42 Å for TM1-TM6 distance distributions. Populations are provided in **Table S3**.

We asked whether there would be differences among AT1R/βarr1* complexes formed in the presence of these β-arrestin-biased ligands. We performed DEER measurements on our TM1-TM6 pair bound to TRV023, TRV026, and TRV055 in the absence and presence of βarr1* **(Fig. 3B)**. TRV055 is a Gq-biased ligand that has enhanced allosteric coupling to Gq and similar coupling to β-arrestin as AngII^29^. Like AngII, it stabilizes the canonical, fully activated AT1R conformation^25^. The addition of βarr1* quantitatively stabilizes the ~25 Å and 42 Å distances for both AngII, TRV023, and TRV055 **(Fig. 3B, 3C)**. However, even though DEER experiments are performed at mid-micromolar protein concentrations, βarr1* does not fully deplete the 35 Å population in the presence of TRV026. The incomplete complex formation observed even at high protein concentrations suggests that βarr1* has substantially lower affinity for TRV026-AT1R than for AngII-AT1R and TRV023-AT1R.

To quantitatively assess the ligand dependence of complex affinity, we employed an AlphaScreen proximity assay that allows us to detect the interaction between FLAG-AT1R and βarr1* (through a His6-tagged antibody fragment) at mid-nanomolar protein concentrations. While TRV023 is at least as effective as AngII in promoting βarr1* complex formation, TRV026 does not give an appreciable signal in this assay **(Fig. 4A)**. Importantly, the difference in βarr1* interaction is not due to lower ligand affinity, since the affinity of TRV026 for purified AT1R is comparable to AngII and higher than TRV023^27^. In addition, TRV055, a much lower affinity ligand^25^, effectively promotes complex formation.

**Fig. 4.**
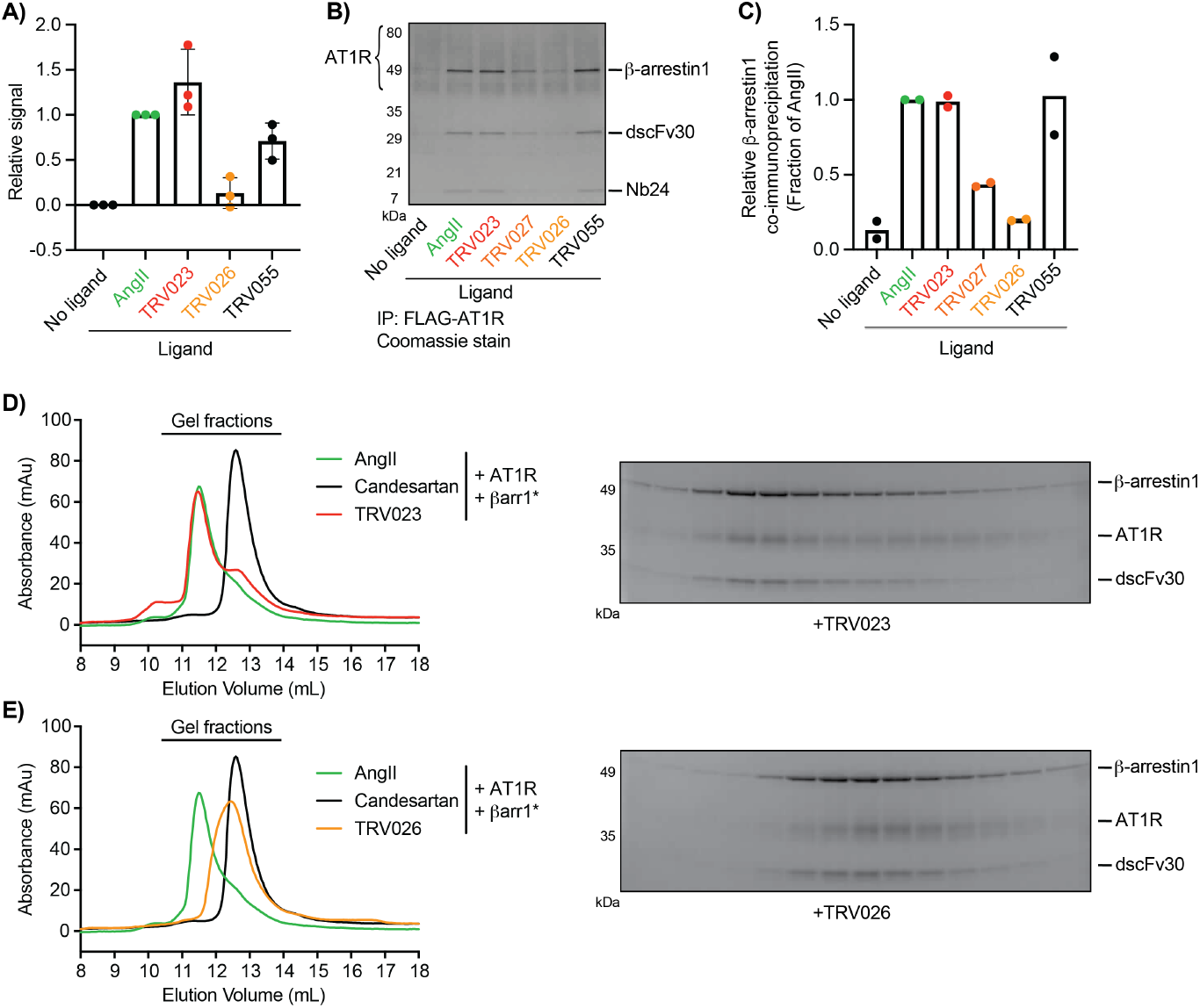
Ligand-dependent stability of AT1R/βarr1* complexes. **(A)** AlphaScreen assay detecting the interaction between FLAG-tagged AT1R (100 nM) and βarr1* (33 nM) (via His6-tagged Fab30) in the presence of saturating concentrations of various ligands. Data are normalized to the no ligand/AngII signals and represent the mean ± S.D. of 3 independent experiments. **(B, C)** Co-immunoprecipitation of βarr1* with AT1R (glycosylated) in the presence of saturating concentrations of various ligands. (B) shows a representative Coomassie-stained gel; (C) shows quantitation of the β-arrestin1 band from 2 independent experiments. **(D**,**E)** Stability of deglycosylated AT1R/βarr1* complexes formed with (D) TRV023 and (E) TRV026 as assessed by SEC and gel electrophoresis of the resulting fractions.

We further asked how various β-arrestin-biased ligands affect the rate of dissociation of AT1R/βarr1* complexes using co-immunoprecipitation. Here complexes are formed at micromolar concentrations, but βarr1* can dissociate during washing. TRV055 and TRV023 promote βarr1* pulldown as effectively as AngII **(Fig. 4B**,**C)**. TRV027 and TRV026 are increasingly less effective.

Finally, we analyzed the ability of biased ligands to stabilize AT1R/βarr1* complexes by SEC. TRV023 complexes remain mostly intact, with the majority eluting at the shorter retention time observed for AngII **(Fig. 4D)**. In contrast, TRV026 complexes largely dissociate, with a broad peak migrating closer to the retention time of candesartan-bound AT1R **(Fig. 4E)**. Together these data demonstrate that βarr1* has lower affinity for the AT1R 7TM core in the presence of TRV026 compared to TRV023 and AngII.

### TRV026-stabilized conformations observed in silico do not accommodate β-arrestin binding to the 7TM core

DEER shows that TRV026 stabilizes a structural intermediate that we previously designated Occluded 1^25^, and our functional data indicate βarr1* cannot effectively engage this conformation through the 7TM core. We used unbiased molecular dynamics (MD) simulations to obtain an all-atom structural model of the TRV026-bound conformation. We performed five MD replicas of 20 μs each to capture conformational transitions between different TM6 conformations that occur on a slow timescale^30^. Simulations were initiated from an inactive conformation of AT1R^31^ (PDB ID: 4YAY), a choice which was motivated by previously reported challenges in capturing large conformational changes in MD simulations initiated from active AT1R^32^. To compare our MD simulations with the experimental DEER distance distributions, we attached spin labels *in silico* to all experimental labeling sites and simulated the corresponding distance distributions for each MD frame (see Materials and Methods). In three out of five replicas, TM6 moved outward by about 2 Å, similar to the distance change observed in DEER (TM1-TM6 distance 32 Å for inactive conformation vs. 35 Å for Occluded 1; **Fig. S2**)^25^.

In one replica, TM6 transitioned after the first 5 µs and stably maintained the intermediate position until the end of the simulation **(Fig. 5A)**. The time evolution of simulated peak positions for all ten spin label pairs in this replica are shown in **Fig. S4**. To determine a representative structure of the intermediate conformation, we calculated the median structure of all MD simulation frames after the first 5 μs **(Fig. 5A)**. This structural model confirms that TM6 assumes an intermediate position relative to inactive- and active-state AT1R structures **(Fig. 5B, Fig. S5)**. Another prominent change is that H8 moves to a membrane-normal orientation. This position of H8 is distinct both from inactive AT1R structures, where it is angled slightly away from the membrane^31^, and from active AT1R structures, where it lies parallel to the membrane^33,34^, as is typical for most GPCRs. A similar non-canonical position of H8 has been observed for other GPCRs, including the angiotensin II type 2 receptor^35,36^. Unlike many GPCRs, AT1R lacks a palmitoylation site in H8, which likely increases structural flexibility in this segment. Notably, our previous DEER data indicated conformational changes in the TRV026-stabilized Occluded 1 conformation that are consistent with the out-of-plane movement of H8^25^.

**Fig. 5.**
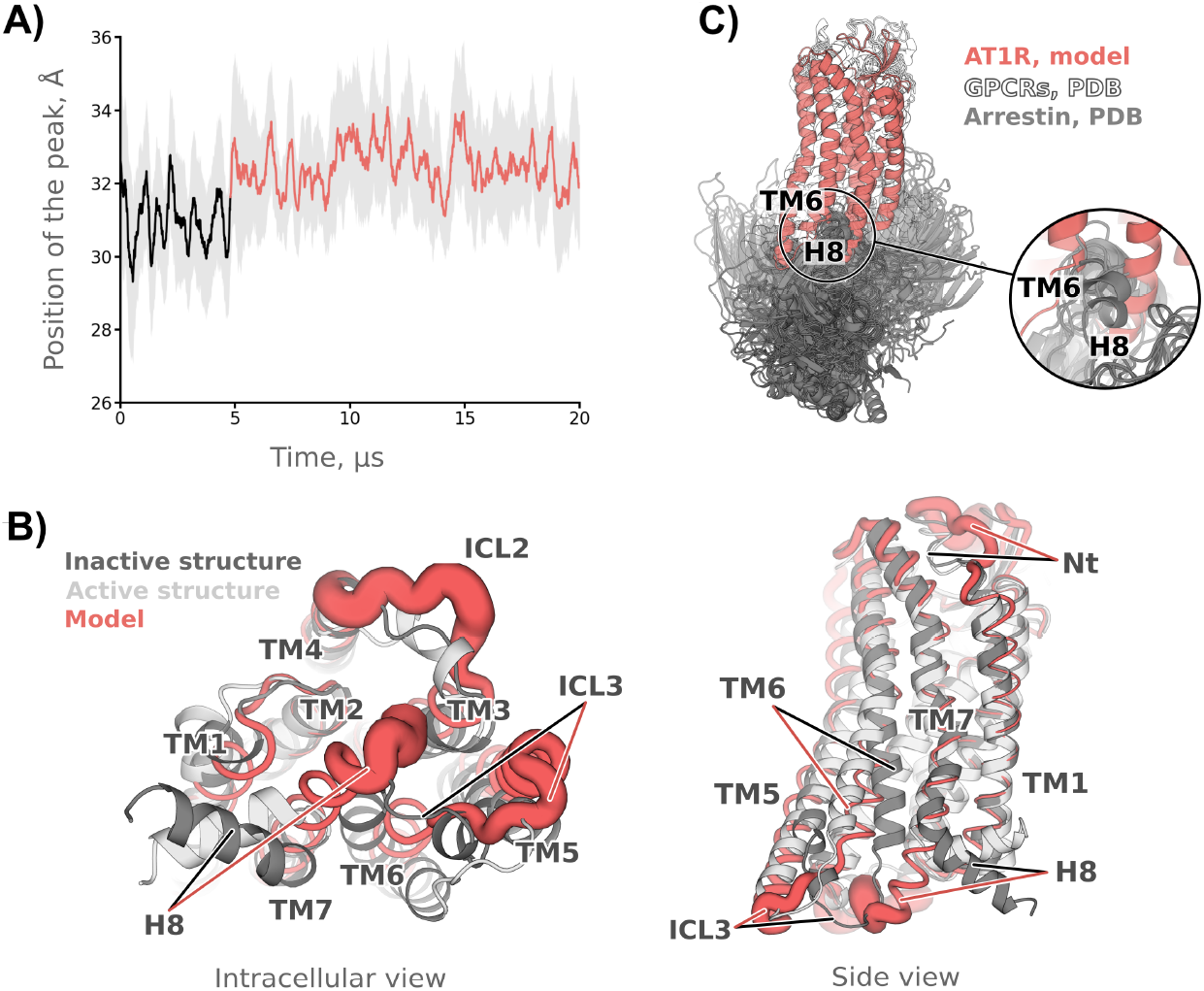
MD analysis of AT1R bound to TRV026. **(A)** Peak distance of the modeled TM1-TM6 distance distribution over simulation time. The bold line represents the sliding mean, and the semi-transparent gray area indicates the standard deviation. After about 5 µs, the inactive-like peak position (black line) shifts toward an intermediate distance (red line). **(B)** Intracellular and side views of the AT1R structural model with the intermediate TM6 position (red). The active (light gray, PDB: 6OS2) and inactive (dark gray, PDB: 4YAY) AT1R structures are shown for comparison. Backbone flexibility is visualized by mapping the RMSF to the width of the cartoon representation. **(C)** Superposition of GPCR–arrestin core complexes with the model of the TRV026-bound intermediate conformation. All observed poses of visual arrestin and β-arrestin1 clash with TM6 and H8 of the AT1R model.

We also evaluated the flexibility of individual protein segments by calculating the backbone root mean square fluctuation (RMSF) values **(Fig. 5B)**. The N-terminus, ICL2 and H8 show the highest flexibility in the intermediate conformation, which may have functional relevance for those segments. The conformational flexibility likely diminishes the accuracy of spin label modeling and calculation of distance distributions for those segments **(Fig. S5)**. ICL3 was less dynamic than those regions, and the 7TM region remained largely stable.

Finally, we asked whether the 7TM core of this TRV026-stabilized AT1R conformation could engage β-arrestins. To this end, we superimposed all available core-engaged, high-resolution structures of GPCRs in complex with β-arrestin1 or visual arrestin on the intermediate model described above **(Fig. 5C)**. Both TM6 and H8 of the intermediate model clash with arrestins, particularly with the finger loop that inserts into the GPCR transducer-binding pocket. This accords with our DEER data showing that driving complex formation between TRV026-bound AT1R and βarr1* at high protein concentrations leads to the outward displacement of TM6 **(Fig. 3B)**.

## Discussion

The multimodal interactions of β-arrestins with GPCRs results in several types of conformational heterogeneity^4,7,37^. First, complexes can sample conformations where β-arrestin is bound only to the phosphosites and where β-arrestin is bound both to the phosphosites and the 7TM core^4,7^. Second, complexes can retain substantial plasticity even when β-arrestin interacts with the 7TM core. In this manuscript, we provide insights into both of these aspects of conformational heterogeneity by developing generalizable methodology to form GPCR/β-arrestin complexes mediated exclusively by 7TM core interactions. Our *in vitro* analysis of AT1R/βarr1* complexes is limited by the detergent environment of the receptor^26,38^ and by the use of a V2R-derived phosphorylated peptide and antibody fragments to stabilize an activated conformation of β-arrestin1. Interactions with membrane lipids^3–6,39,40^ and different phosphorylation patterns^41,42^ could influence the conformational distribution of β-arrestin and its interaction with the AT1R 7TM core. Nevertheless, our findings suggest a potentially important mechanism through which ligands could bias the activation of different β-arrestin-mediated pathways.

Numerous biophysical approaches have revealed conformational heterogeneity in complexes where β-arrestin is fully engaged with the GPCR 7TM core. High-resolution cryo-electron microscopy structures show that β-arrestin can assume many orientations relative to the 7TM core of various GPCRs^4,5,37,43–46^. Intermolecular DEER measurements of rhodopsin/visual arrestin^47^ and crosslinking studies of the parathyroid hormone 1 receptor/β-arrestin1 in live cells^48^ provide evidence of flexibility within these complexes. NMR studies of GPCRs bound to mutationally activated β-arrestin1 have highlighted the dynamics of residues throughout the receptors, from the ligand-binding site to the intracellular face^49,50^. Consistent with these studies, our DEER data show conformational heterogeneity in certain intracellular regions of the AT1R, particularly H8, when bound to βarr1* **(Fig. 2B)**. On the other hand, several intracellular receptor segments, such as TM6, TM7 and ICL2, are stabilized by βarr1*.

AngII, TRV023, and TRV026 each stabilize distinct AT1R conformational distributions in the absence of transducers^25^. Yet to the extent that our DEER probes can resolve, the same intracellular conformation dominates in the presence of high concentrations of βarr1*, regardless of which ligand is bound **(Fig. 3B)**. This is not surprising since the intracellular AT1R conformation in the final complex will be largely dictated by the direct interactions this region makes with the transducer^27,51^. In contrast, the addition of β-arrestin1 to phosphorylated μ-opioid receptor (μOR) bound to full, partial, or biased agonists had minimal effects on the μOR intracellular conformational distribution by DEER^52^. The difference between the AT1R and μOR systems may be due to the relatively low affinity of β-arrestin1 for the μOR 7TM core. We previously demonstrated that μOR/β-arrestin1 complexes are predominantly in conformations where β-arrestin1 only engages the phosphosites^53^.

Despite the similarity of the dominant AT1R conformation in βarr1* complexes formed with various orthosteric ligands, we found that βarr1* has a lower affinity for and dissociates more quickly from AT1R bound to TRV026 **(Fig. 4)**. TRV026 stabilizes the Occluded 1 conformation that is characterized by an intermediate position of TM6. Ligands that stabilize conformations with a fully displaced TM6 promote more stable complex formation. This includes both TRV023, which stabilizes the Occluded 2 conformation, and ligands such as AngII and TRV055, which stabilize the canonical, fully activated conformation^25^. This indicates that even though the Occluded 1 and Occluded 2 conformations are both stabilized by β-arrestin-biased AT1R ligands, they have different efficacies toward βarr1* engagement of the core. Previous data have shown that TRV027 only promotes interactions of β-arrestins with the phosphosites, while AngII promotes engagement of both the phosphosites and 7TM core^54^. This results from these ligands’ differential activation of Gq and GRKs^54,55^. Our data suggest that even when ligands have similar bias toward transducers other than β-arrestin, the allosteric effects of orthosteric ligands on the 7TM core could also directly regulate the conformations of GPCR/β-arrestin complexes.

Even though TRV026, TRV027, and TRV023 differ in their ability to stabilize the 7TM core interaction, they have similar efficacies and potencies in assays of β-arrestin recruitment, internalization, and kinase pathway activation^27,28^ **(Fig. 3A)**. All these functional responses can be mediated by β-arrestins when they are restricted to “phosphosites only” conformations, in the absence of 7TM core engagement^8^. Therefore, these assays likely reflect the comparable ability of these β-arrestin-biased ligands to promote GRK phosphorylation of the C-terminus, the main driver of β-arrestin recruitment to the AT1R^54,56^. Other responses mediated by β-arrestins, such as desensitization of G protein activation by steric blockade, are known to correlate with the strength of the 7TM core interaction^8,53^. In some systems, the core interaction has also been shown to modulate ERK activation^10^. Altering the equilibrium between the “phosphosites” and “phosphosites + core” conformations thus provides a potential mechanism to bias the manifold functions of β-arrestins. Notably, several β-arrestin-biased AT1R ligands have shown qualitative differences in both proximal β-arrestin activation assays and downstream signaling readouts, providing further evidence of intra-transducer bias within this pharmacological class^57,58^.

Our MD simulations provide a molecular rationale for the inability of βarr1* to engage the 7TM core of TRV026-bound AT1R effectively **(Fig. 5)**, despite the structural plasticity of arrestins. Conversely, the increased dynamics of H8 observed by both DEER^25^ and in simulation **(Fig. 5B)** upon TRV026 binding could facilitate the exposure of C-terminal sites for phosphorylation by GRKs. The ligand dependence of β-arrestin interactions with the 7TM core may vary among GPCRs. NMR studies indicate that β-arrestin1 in complex with a phosphorylated β2-adrenergic receptor (β2AR)/V2R chimera undergoes ligand-dependent conformational changes that do not consistently correlate with ligands’ efficacy towards G protein activation or β-arrestin-dependent internalization^59^. This suggests that β-arrestin is sensitive to the conformation of the β2AR core, as we have shown here for AT1R. In contrast, purified β1-adrenergic receptor forms 7TM-core-mediated complexes with V2Rpp-activated β-arrestin1 equally well regardless of whether no ligand, an antagonist, an agonist, or a β-arrestin-biased ligand is bound^60^.

In summary, our data show that the different conformations stabilized by β-arrestin-biased AT1R ligands determine their ability to promote β-arrestin interactions with the 7TM core. There are now several examples of functionally similar ligands that activate the same GPCR through distinct mechanisms, including M2 muscarinic receptor partial agonists^61^ and β2AR partial agonists^62^. This raises the possibility that there may be other finer levels of ligand bias—and additional opportunities to tune the pharmacology of GPCR drugs—that would not be detected in standard high-throughput screening assays. Most β-arrestin activation assays monitor its recruitment to GPCRs, principally reflecting β-arrestin’s high-affinity interaction with GRK-phosphorylated regions. On the other hand, efforts to rationally design β-arrestin-biased ligands have been based on 7TM core-engaged GPCR/β-arrestin structures. Our findings suggest that these interactions may not always be correlated. To maximize the likelihood of developing GPCR drugs with the desired pharmacological profiles, it will be important to determine how different modes of β-arrestin interaction influence the cellular responses of interest and to tailor drug discovery approaches accordingly.

## Methods

### Cell lines

Expi293F Inducible cells (Thermo) were grown in Expi293 expression media (Thermo) with shaking at 110 rpm in a humidified 37 °C incubator with 8% CO_2_.

### Protein expression and purification

FLAG-tagged AT1R constructs were expressed in Expi293F Inducible cells transiently transfected with the Expifectamine transfection kit (Thermo) as previously described^25,33^, and expression was induced two days after transfection with 4 μg/mL doxycycline and 5 mM sodium butyrate in the presence of 1 um losartan, a low-affinity antagonist. Cells were harvested approximately 30 hours after induction, washed with phosphate-buffered saline and 1 μM losartan, and flash frozen. Cells were lysed under hypotonic conditions (10 mM Tris pH 7.4, 2 mM EDTA, 10 mM MgCl_2_, 1 μM losartan, 2.5 U/mL benzonase (Sigma), protease inhibitors benzamidine and leupeptin) and then dounced and solubilized for 2 hours at 4 °C (0.5% lauryl maltose neopentyl glycol (MNG), 0.05% cholesterol hemisuccinate (CHS), 20 mM HEPES pH 7.4, 500 mM NaCl, 10 mM MgCl_2_, 1 μM losartan, benzonase, benzamidine, leupeptin). After centrifugation (30,000 × g, 30 min, 4 °C), 2 mM CaCl_2_ was added, and solubilized material was loaded onto M1 anti-FLAG resin at 4 °C. The resin was washed (20 mM HEPES pH 7.4, 500 mM NaCl, 0.01% MNG, 0.01% CHS, 2 mM CaCl_2_, benzamidine, leupeptin) and eluted (20 mM HEPES pH 7.4, 500 mM NaCl, 0.01% MNG, 0.01% CHS, 0.2 mg/mL FLAG-peptide). The protein was subjected to size exclusion chromatography (SEC) (Superdex 200 Increase 10/300 GL column, GE) in buffer consisting of 20 mM HEPES pH 7.4, 100 mM NaCl, 0.01% MNG, 0.001% CHS, and monomeric fractions were concentrated.

For DEER experiments, AT1R was eluted from M1 resin using buffer at pH 6.8 and incubated with a 20-fold molar excess of bis-(2,2,5,5-tetramethyl-3-imidazoline-1-oxyl-4-yl)disulfide (Enzo) for 3 hours at room temperature. SEC buffer was made with D_2_O and contained 40 mM HEPES. For AT1R used in analytical SEC experiments, transfected cells were cultured in the presence of 5 uM kifunensine, and M1-purified receptor was deglycosylated with EndoH for 90 minutes at room temperature before the SEC purification step^33^.

Cysteine-free rat β-arrestin1 truncated at residue 393 was purified as a thrombin-cleavable GST fusion from BL21 DE3 *E. coli* as previously described^53^. Briefly, cells were lysed with a microfluidizer, and the fusion was captured on glutathione sepharose resin. After thrombin digest, β-arrestin1 was isolated by ion exchange chromatography (HiTrap Q sepharose). His-tagged antibody fragments that recognize an activated conformation of β-arrestin1 were purified as follows. Fab30 was purified as described previously^23^. A disulfide-stabilized single chain version of Fab30 (dscFv30)^4^ and the nanobody Nb24^4^ were expressed in T7 Express lysY BL21 *E. coli* (New England Biolabs) and purified from the periplasmic fraction via NiNTA chromatography as previously described^33^; dscFv30 was expressed as a 3C-protease cleavable maltose-binding protein (MBP) fusion. To form the βarr1* complex, β-arrestin1 was incubated overnight at 4 °C with a 1.25 molar excess of V2Rpp phosphopeptide (ARGRpTPPpSLGPQDEpSCpTpTApSpSpSLAKDTSS), MBP-dscFv30, and Nb24. The complex was treated with 3C protease for 90 minutes at room temperature and purified by SEC (Superdex 200 Increase 10/300 GL column) in 20 mM HEPES pH 7.4, 100 mM NaCl. For DEER experiments, SEC buffer was made with D_2_O.

### DEER experiments

DEER samples consisted of spin-labeled AT1R (110 μM), βarr1* (140 μM), 1 mM ligand, and 20% d^8^-glycerol (Cambridge Isotopes) in a final volume of 12 μL. After a 1 hour incubation at room temperature, binding reactions were transferred to in 1.4/1.7 mm (i.d./o.d.) borosilicate capillaries (VitroCom, Mountain Lakes, NJ), flash frozen, and stored at −80 °C. 4-Pulse DEER experiments were conducted at 50 K on a Q-band upgraded ElexSys 580 (Bruker, Germany) equipped with an arbitrary waveform generator, a 150 W traveling wave tube amplifier and a QT-2 probehead (Bruker) using the standard dead time free sequence^63^. Pulse lengths were optimized using echo nutation experiments and set to 16-20 ns and 32-40 ns for π/2 and π-pulses, respectively. A 100 ns 50 MHz-chirp pulse was applied as a pump pulse 70 MHz above the observer frequency. All data, including the full list of pulse parameters, are accessible via https://doi.org/10.5281/zenodo.15590546.

### DEER data analysis

Dipolar evolutions were recorded with 16 ns time resolution and analyzed model-free in LongDistances v.1112^64^. The smoothness parameter was determined by L-curve criterion. Standard deviations for distances were determined using the error analysis implementation in LongDistances using standard parameters and the inherited model-free parameters. Population analysis was performed using a global Gaussian mixture model implemented in the DeerLab Python toolbox (v.1.0.2). Positions and widths of the Gaussians were treated as global fitting parameters, while populations, concentrations and modulation depths were local parameters. 95% confidence bands were estimated via bootstrapping (100 iterations).

### Radioligand binding

Purified AT1R (75 ng) was incubated with βarr1* (2.5 μM), [^3^H]-olmesartan (2.5 nM) (American Radiolabeled Chemicals) and varying concentrations of cold competitor ligand in assay buffer (20 mM HEPES pH 7.4, 100 mM NaCl, 0.01% MNG, 0.001% CHS, 0.1% BSA) in a total volume of 200 μL for 2 hours at room temperature. Binding reactions were harvested onto GF/B filters using a 96-well Brandel harvester and washed with cold 20 mM HEPES, 100 mM NaCl. Bound radioactivity was measured using a Packard TriCarb 2100TR scintillation counter, and a 3-parameter dose-response curve was fit to the normalized data from 3 independent experiments (GraphPad Prism).

### Analytical SEC

Deglycosylated AT1R (12 μM) was incubated with ligand (333 μM) for 15 minutes at room temperature, and βarr1* (12 μM) was added for a total volume of 100 μL. After an additional 1 hour incubation at room temperature, samples were loaded onto SEC (Superdex 200 Increase 10/300 GL column, 0.5 mL/min flow rate, 0.25 mL fractions). Fractions were analyzed by gel electrophoresis.

### Co-immunoprecipitation

Purified AT1R (7 μM) was incubated for 1 hour at room temperature with ligand (1 mM) and βarr1* (9 μM) in assay buffer (20 mM HEPES pH 7.4, 100 mM NaCl, 0.001% MNG, 0.0001% CHS). Additional assay buffer (30 μL), CaCl_2_ (2 mM) and M1 anti-FLAG resin (20 μL) were added, and samples were rotated for 20 minutes at room temperature. The resin was washed 4 times with assay buffer containing 2 mM CaCl_2_ and then eluted with assay buffer containing 5 mM EDTA and FLAG peptide. Eluates were analyzed by gel electrophoresis, and bands were quantified in ImageJ.

### AlphaScreen

βArr1* complex was formed *in situ* by incubating His-tagged Fab30 (17 μM) with a 2-fold excess of β-arrestin1 and V2Rpp for 1 hour at room temperature and then diluting 250-fold in 20 mM HEPES pH 7.4, 100 mM NaCl, 0.1% BSA for a working Fab30 concentration of 67 nM. Purified AT1R (200 nM) was incubated with ligand (200 μM) for 15 minutes at room temperature in 20 mM HEPES pH 7.4, 100 mM NaCl, 0.01% MNG/0.001% CHS, 0.1% BSA. βarr1* (10 μL) and ligand-bound AT1R (10 μL) were combined and incubated for 1 hour before the simultaneous addition of anti-FLAG-conjugated AlphaScreen Acceptor beads (0.05 mg/well) and Nickel Chelate Alpha Donor beads (0.05 mg/well) (Perkin Elmer) in a 5 μL volume. The signal resulting from the close proximity of FLAG-AT1R and Fab30-His6 was read on a CLARIOstar plate reader (BMG Labtech) after 1 hour.

### β-arrestin recruitment assay

The LgBit-CAAX and β-arrestin1-SmBit plasmids were a gift from Sudar Rajagopal (Duke University). Inducible Expi293 cells were co-transfected with LgBit-CAAX, β-arrestin2-SmBit, and pcDNA-Zeo-tetO-Flag-AT1R^33^ in a 1:4:5 ratio. Cells were induced 2 days post-transfection as described above. Three days post-transfection, cells were harvested and resuspended in Hanks’ balanced salt solution + 20 mM HEPES pH 7.4 (assay buffer) containing Coelenterazine H (Research Products International) (3.1 μM), and 80 μL of cells were dispensed into opaque white 96-well plates. After reading basal luminescence 3 times (Clariostar, BMG LabTech), cells were treated with 20 μL of a serial dilution of ligands prepared in assay buffer containing 0.1% BSA. Luminescence values collected 20 minutes post-stimulation were normalized to the average of the three basal luminescence reads from the same well. A 3-parameter dose-response curve was fit to the normalized data from 3 independent experiments (GraphPad Prism).

### Preparation of model for MD simulations

An X-ray crystal structure of the AT1R in the inactive conformation (PDB: 4YAY) was used as a starting structure for the MD simulations^31^. The co-crystallized small-molecule antagonist ZD7155 and the soluble cytochrome b562 were removed. The missing regions of the ECL2 and ICL3 were reconstructed using AlphaFold2^65^. The refined protein structure was aligned to the TM1, 2 and 4 helices of the entry 4YAY from the Orientation of Proteins in Membranes (OPM) database^66^. Next, TRV026 was placed into the orthosteric ligand binding pocket of the reconstructed inactive conformation using the ligand conformation and orientation from the TRV026-bound active AT1R structure (PDB: 6OS2)^27^. The simulation box of the system including water molecules, ions, lipids, and the TRV026-bound AT1R was built using CHARMM-GUI^67^. All residues were kept in the standard protonation states of the CHARMM36m force field, except D74^2.50^ and D125^3.49^, which were set as protonated. AT1R, in contrast to many class A GPCR representatives, does not have a sodium-binding site in proximity to D74^2.50^, and, in the absence of sodium, D74^2.50^ is likely to be protonated^32^. D125^3.49^ is normally deprotonated in the inactive conformation and protonated in the active conformation; thus, we set D125 as protonated in order to drive our simulation away from the inactive conformation^25^. The ligand-bound receptor was oriented in the membrane with the “Use PDB Orientation” option of the CHARMM-GUI. The number of POPC lipids in each upper- and lower leaflet was defined as 75. Na^+^ and Cl^−^ ions were added to neutralize the net charge and reach physiological ionic strength, 0.15 M. The resulting output files with coordinates, topology, atom indexes, and parameters of the CHARMM36m force field were downloaded, and force field parameters for the non-standard residue Sar1 of TRV026 were adopted from the earlier publication^32^. The resulting system contained 70,200 atoms, 14,902 TIP3 water molecules, 150 POPC lipids, 39 Na^+^ and 58 Cl^-^ ions. The rectangular simulation box was 79.062 × 79.062 × 12.082 Å.

### MD simulations

We performed five independent replicas of unbiased molecular dynamics (MD) simulations using GROMACS, version 2021.3^68^. Each starting system was first subjected to energy minimization using 10,000 steps of the steepest descent algorithm. During minimization, both positional (POSRE) and dihedral (DIHRES) restraints were applied with the following force constants (in kJ mol^1^ nm^2^ for POSRE and kJ mol^1^ rad^2^ for DIHRES):

|  |  |
| --- | --- |
| POSRE backbone | 4000.0 |
| POSRE side chains | 2000.0 |
| POSRE lipids | 1000.0 |
| DIHRES | 1000.0 |

The equilibration phase consisted of six sequential steps. The first was a 125 ps NVT equilibration using the same force constants as during energy minimization. The second step was also 125 ps of NVT equilibration but with the reduced values of force constants (see table below). All equilibration simulations were conducted at a target temperature of 303.15 K using the Berendsen thermostat^69^. The following four steps were NPT equilibration simulations: 125 ps in the third step, 500 ps in fourth step, 20,000 ps in fifth step, and 500 ps in sixth step. The restraint strengths were progressively reduced across the steps as follows:

|  | step 2 | step 3 | step 4 | step 5 | step 6 |
| --- | --- | --- | --- | --- | --- |
| POSRE backbone | 2000.0 | 1000.0 | 500.0 | 200.0 | 50.0 |
| POSRE side chains | 1000.0 | 500.0 | 200.0 | 50.0 | 0.0 |
| POSRE lipids | 400.0 | 400.0 | 200.0 | 40.0 | 0.0 |
| DIHRES | 400.0 | 200.0 | 200.0 | 100.0 | 0.0 |

During equilibration, pressure was controlled using a semi-isotropic Berendsen barostat, with the xy-plane and z-axis coupled independently^69^. A time step of 1 fs was used for equilibration steps 1–3, and 2 fs for steps 4–6. The overall equilibration time was 21,250 ps.

Each replica was followed by a 20 μs production MD simulation, carried out without any positional or dihedral constraints. Temperature and pressure were maintained using the velocity-rescale thermostat and the semi-isotropic Parrinello–Rahman barostat, respectively^70,71^. Long-range electrostatics were handled using the Particle Mesh Ewald (PME) method, with fourth-order interpolation and a real-space cut-off of 1.2 nm^72,73^. Van der Waals interactions were switched off smoothly between 1.0 and 1.2 nm. All covalent bonds involving hydrogen atoms were constrained to their equilibrium lengths using P-LINCS^74^. A 2 fs integration time step was used during production, with Verlet neighbor lists updated every 20 fs and the neighbor list radius set automatically.

### Preparation of MD trajectory for analysis

For the analysis, we reconstructed periodic boundary conditions (PBC) using “gmx trjconv” command to ensure that the entire molecular system remained within a single unit cell without broken covalent bonds due to periodicity. We then applied procedures of rotation and translation to align AT1R to its starting structure using “gmx trjconv”. The alignment was performed on the backbone heavy atoms of TM1, TM2, and TM4, since these helices are known to remain relatively rigid across different conformational states of GPCRs^75,76^. This selection of atoms for alignment allowed us to minimize alignment artifacts. This postprocessing protocol eliminated the need for additional alignment steps in subsequent analyses and enabled more straightforward visual inspection of the trajectories using PyMOL.

To reduce computational complexity for downstream analyses, we generated a shortened version of the 20 μs-long post-processed MD trajectory. This reduced dataset consisted of 2000 frames, extracted at 10 ns intervals.

### Measurement of TM1–TM6 distance

In this study, the “TM1–TM6” distance refers to the distance between the Cα atoms of residues F55^1.59^ and D236^6.31^. We calculated this distance along the MD trajectory using the MDanalysis Python package^77,78^. We plotted changes of the TM1–TM6 distance across the time-trace as a semi-transparent line **(Fig. 5A)**, with a bold line of a sliding mean.

### Simulation of DEER distance distributions for an MD trajectory

To simulate DEER distance distributions, we used the chiLife Python package^79^. Typically, chiLife works with single structures loaded into a Universe object. Since the Universe object is a feature of the MDAnalysis, we utilized this library to load the entire MD trajectory into the Universe object. This enabled us to iterate through the MD simulation frames and simulate DEER distance distributions for each frame individually.

For our analysis, we used the V1 spin label, consistent with that used in the corresponding DEER experiments^25^. The labeling positions are shown in **Fig. 2A**. At each labeling site, ensembles of V1 spin label conformers were generated using the Accessible Volume (AV) approach, with 10,000 iterations^79^.

### Calculation of RMSF

We computed Root Mean Square Fluctuation (RMSF) for all Cα atoms of AT1R. To do this, we used MD simulation frames recorded after 5 μs, as these represented the intermediate structure.

The RMSF values were added to the B-factor column of the .pdb file with the representative intermediate structure. To illustrate flexibility of different segments of AT1R, we visualized the obtained RMSF values as a putty plot **(Fig. 5B)**, using PyMOL version 2.5 Schrödinger, LLC^80^.

### Alignment of the model with GPCR-arrestin complexes

To investigate why arrestin binding to TRV026-bound AT1R may be hindered, we aligned GPCR–arrestin complexes from the PDB to the structural model of the intermediate conformation. All downloaded GPCR–arrestin complexes were aligned in PyMol using the backbone heavy atoms. The PDB identifiers of the GPCR– arrestin complexes used in the alignment are listed in **Table S4**.

## Data, Materials, and Software Availability

Raw DEER data are deposited at https://doi.org/10.5281/zenodo.15590546. All other data are included in the manuscript and/or the supplemental information.

## ACKNOWLEDGMENTS

We thank Robert J. Lefkowitz for critical reading of this manuscript. We thank Wayne L. Hubbell for the use of his equipment for DEER experiments. The authors gratefully acknowledge the computing time provided by them on the high-performance computers Barnard and Romeo at NHR@TUD, which is funded by the Federal Ministry of Education and Research and the state governments participating on the basis of the resolutions of the GWK for the national high-performance computing at universities (www.nhr-verein.de/unsere-partner). This work was funded by the German Federal Ministry of Education and Research, SECAI 57616814 to J.B., and the German Research Foundation, project number 421152132, subproject A03 and project number 514664767, subproject B07 to M.E. L.M.W. is a Whitehead Scholar and a Pew Scholar in the Biomedical Sciences, supported by The Pew Charitable Trusts.

## AUTHOR CONTRIBUTIONS

M.E. and L.M.W. conceived the study. D.P.S. contributed new reagents and protocols. L.M.W. purified proteins, prepared DEER samples, and performed *in vitro* biochemical experiments, radioligand binding, and cellular assays. M.E. performed DEER experiments and data analysis. M.B.H. and J.B. performed and analyzed MD simulations. J.B., M.E., and L.M.W. wrote the manuscript.

## References

1. Gurevich, V.V., Gurevich, E.V., and Uversky, V.N. (2018). Arrestins: structural disorder creates rich functionality. Protein Cell 9, 986–1003. 10.1007/s13238-017-0501-8.

2. Haider, R.S., Reichel, M., Matthees, E.S.F., and Hoffmann, C. (2023). Conformational flexibility of beta-arrestins - How these scaffolding proteins guide and transform the functionality of GPCRs. Bioessays 45, e2300053. 10.1002/bies.202300053.

3. Chen, K., Zhang, C., Lin, S., Yan, X., Cai, H., Yi, C., Ma, L., Chu, X., Liu, Y., Zhu, Y., et al. (2023). Tail engagement of arrestin at the glucagon receptor. Nature 620, 904–910. 10.1038/s41586-023-06420-x.

4. Staus, D.P., Hu, H., Robertson, M.J., Kleinhenz, A.L.W., Wingler, L.M., Capel, W.D., Latorraca, N.R., Lefkowitz, R.J., and Skiniotis, G. (2020). Structure of the M2 muscarinic receptor-beta-arrestin complex in a lipid nanodisc. Nature 579, 297–302. 10.1038/s41586-020-1954-0.

5. Huang, W., Masureel, M., Qu, Q., Janetzko, J., Inoue, A., Kato, H.E., Robertson, M.J., Nguyen, K.C., Glenn, J.S., Skiniotis, G., and Kobilka, B.K. (2020). Structure of the neurotensin receptor 1 in complex with beta-arrestin 1. Nature 579, 303–308. 10.1038/s41586-020-1953-1.

6. Chen, Q., Schafer, C.T., Mukherjee, S., Wang, K., Gustavsson, M., Fuller, J.R., Tepper, K., Lamme, T.D., Aydin, Y., Agrawal, P., et al. (2025). Effect of phosphorylation barcodes on arrestin binding to a chemokine receptor. Nature 643, 280–287. 10.1038/s41586-025-09024-9.

7. Shukla, A.K., Westfield, G.H., Xiao, K., Reis, R.I., Huang, L.Y., Tripathi-Shukla, P., Qian, J., Li, S., Blanc, A., Oleskie, A.N., et al. (2014). Visualization of arrestin recruitment by a G-protein-coupled receptor. Nature 512, 218–222. 10.1038/nature13430.

8. Cahill, T.J., 3rd, Thomsen, A.R., Tarrasch, J.T., Plouffe, B., Nguyen, A.H., Yang, F., Huang, L.Y., Kahsai, A.W., Bassoni, D.L., Gavino, B.J., et al. (2017). Distinct conformations of GPCR-beta-arrestin complexes mediate desensitization, signaling, and endocytosis. Proc Natl Acad Sci U S A 114, 2562–2567. 10.1073/pnas.1701529114.

9. Kumari, P., Srivastava, A., Banerjee, R., Ghosh, E., Gupta, P., Ranjan, R., Chen, X., Gupta, B., Gupta, C., Jaiman, D., and Shukla, A.K. (2016). Functional competence of a partially engaged GPCR-beta-arrestin complex. Nat. Commun. 7, 13416. 10.1038/ncomms13416.

10. Matthees, E.S.F., Haider, R.S., Klement, L., Reichel, M., Blum, N.K., Weitzel, V., Trupschuch, T., Ziegler, C., Drube, J., Schulz, S., and Hoffmann, C. (2025). Helix-bundle and C-terminal GPCR domains differentially influence GRK-specific functions and beta-arrestin-mediated regulation. Nat. Commun. 16, 5430. 10.1038/s41467-025-61281-4.

11. Rankovic, Z., Brust, T.F., and Bohn, L.M. (2016). Biased agonism: An emerging paradigm in GPCR drug discovery. Bioorg Med Chem Lett 26, 241–250. 10.1016/j.bmcl.2015.12.024.

12. Karnik, S.S., Unal, H., Kemp, J.R., Tirupula, K.C., Eguchi, S., Vanderheyden, P.M., and Thomas, W.G. (2015). International Union of Basic and Clinical Pharmacology. XCIX. Angiotensin Receptors: Interpreters of Pathophysiological Angiotensinergic Stimuli. Pharmacol. Rev. 67, 754–819. 10.1124/pr.114.010454.

13. Liu, S., Anderson, P.J., Rajagopal, S., Lefkowitz, R.J., and Rockman, H.A. (2024). G Protein-Coupled Receptors: A Century of Research and Discovery. Circ. Res. 135, 174–197. 10.1161/CIRCRESAHA.124.323067.

14. Abraham, D.M., Davis, R.T., 3rd, Warren, C.M., Mao, L., Wolska, B.M., Solaro, R.J., and Rockman, H.A. (2016). beta-Arrestin mediates the Frank-Starling mechanism of cardiac contractility. Proc. Natl. Acad. Sci. U.S.A. 113, 14426–14431. 10.1073/pnas.1609308113.

15. Kim, K.S., Abraham, D., Williams, B., Violin, J.D., Mao, L., and Rockman, H.A. (2012). beta-Arrestin-biased AT1R stimulation promotes cell survival during acute cardiac injury. Am. J. Physiol. Heart Circ. Physiol. 303, H1001–1010. 10.1152/ajpheart.00475.2012.

16. Cotter, G., Davison, B.A., Butler, J., Collins, S.P., Ezekowitz, J.A., Felker, G.M., Filippatos, G., Levy, P.D., Metra, M., Ponikowski, P., et al. (2018). Relationship between baseline systolic blood pressure and long-term outcomes in acute heart failure patients treated with TRV027: an exploratory subgroup analysis of BLAST-AHF. Clin. Res. Cardiol. 107, 170–181. 10.1007/s00392-017-1168-0.

17. Pang, P.S., Butler, J., Collins, S.P., Cotter, G., Davison, B.A., Ezekowitz, J.A., Filippatos, G., Levy, P.D., Metra, M., Ponikowski, P., et al. (2017). Biased ligand of the angiotensin II type 1 receptor in patients with acute heart failure: a randomized, double-blind, placebo-controlled, phase IIB, dose ranging trial (BLAST-AHF). Eur. Heart J. 38, 2364–2373. 10.1093/eurheartj/ehx196.

18. Self, W.H., Shotwell, M.S., Gibbs, K.W., de Wit, M., Files, D.C., Harkins, M., Hudock, K.M., Merck, L.H., Moskowitz, A., Apodaca, K.D., et al. (2023). Renin-Angiotensin System Modulation With Synthetic Angiotensin (1-7) and Angiotensin II Type 1 Receptor-Biased Ligand in Adults With COVID-19: Two Randomized Clinical Trials. JAMA 329, 1170–1182. 10.1001/jama.2023.3546.

19. Ryba, D.M., Li, J., Cowan, C.L., Russell, B., Wolska, B.M., and Solaro, R.J. (2017). Long-Term Biased beta-Arrestin Signaling Improves Cardiac Structure and Function in Dilated Cardiomyopathy. Circulation 135, 1056–1070. 10.1161/CIRCULATIONAHA.116.024482.

20. Jara, Z.P., Harford, T.J., Singh, K.D., Desnoyer, R., Kumar, A., Srinivasan, D., and Karnik, S.S. (2023). Distinct Mechanisms of beta-Arrestin-Biased Agonist and Blocker of AT1R in Preventing Aortic Aneurysm and Associated Mortality. Hypertension 80, 385–402. 10.1161/HYPERTENSIONAHA.122.19232.

21. Von Moo, E., Harpsoe, K., Hauser, A.S., Masuho, I., Brauner-Osborne, H., Gloriam, D.E., and Martemyanov, K.A. (2022). Ligand-directed bias of G protein signaling at the dopamine D(2) receptor. Cell Chem. Biol. 29, 226–238 e224. 10.1016/j.chembiol.2021.07.004.

22. Motso, A., Pelcman, B., Kalinovich, A., Kahlous, N.A., Bokhari, M.H., Dehvari, N., Halleskog, C., Waara, E., de Jong, J., Cheesman, E., et al. (2025). GRK-biased adrenergic agonists for the treatment of type 2 diabetes and obesity. Cell. 10.1016/j.cell.2025.05.042.

23. Shukla, A.K., Manglik, A., Kruse, A.C., Xiao, K., Reis, R.I., Tseng, W.C., Staus, D.P., Hilger, D., Uysal, S., Huang, L.Y., et al. (2013). Structure of active beta-arrestin-1 bound to a G-protein-coupled receptor phosphopeptide. Nature 497, 137–141. 10.1038/nature12120.

24. Xiao, K., Shenoy, S.K., Nobles, K., and Lefkowitz, R.J. (2004). Activation-dependent conformational changes in beta-arrestin 2. J. Biol. Chem. 279, 55744–55753. 10.1074/jbc.M409785200.

25. Wingler, L.M., Elgeti, M., Hilger, D., Latorraca, N.R., Lerch, M.T., Staus, D.P., Dror, R.O., Kobilka, B.K., Hubbell, W.L., and Lefkowitz, R.J. (2019). Angiotensin Analogs with Divergent Bias Stabilize Distinct Receptor Conformations. Cell 176, 468–478 e411. 10.1016/j.cell.2018.12.005.

26. Van Eps, N., Caro, L.N., Morizumi, T., Kusnetzow, A.K., Szczepek, M., Hofmann, K.P., Bayburt, T.H., Sligar, S.G., Ernst, O.P., and Hubbell, W.L. (2017). Conformational equilibria of light-activated rhodopsin in nanodiscs. Proc. Natl. Acad. Sci. U.S.A 114, E3268–E3275. 10.1073/pnas.1620405114.

27. Wingler, L.M., Skiba, M.A., McMahon, C., Staus, D.P., Kleinhenz, A.L.W., Suomivuori, C.M., Latorraca, N.R., Dror, R.O., Lefkowitz, R.J., and Kruse, A.C. (2020). Angiotensin and biased analogs induce structurally distinct active conformations within a GPCR. Science 367, 888–892. 10.1126/science.aay9813.

28. Violin, J.D., DeWire, S.M., Yamashita, D., Rominger, D.H., Nguyen, L., Schiller, K., Whalen, E.J., Gowen, M., and Lark, M.W. (2010). Selectively engaging beta-arrestins at the angiotensin II type 1 receptor reduces blood pressure and increases cardiac performance. J. Pharmacol. Exp. Ther. 335, 572–579. 10.1124/jpet.110.173005.

29. Strachan, R.T., Sun, J.P., Rominger, D.H., Violin, J.D., Ahn, S., Rojas Bie Thomsen, A., Zhu, X., Kleist, A., Costa, T., and Lefkowitz, R.J. (2014). Divergent transducer-specific molecular efficacies generate biased agonism at a G protein-coupled receptor (GPCR). J. Biol. Chem. 289, 14211–14224. 10.1074/jbc.M114.548131.

30. Dror, R.O., Arlow, D.H., Maragakis, P., Mildorf, T.J., Pan, A.C., Xu, H., Borhani, D.W., and Shaw, D.E. (2011). Activation mechanism of the beta2-adrenergic receptor. Proc. Natl. Acad. Sci. U.S.A 108, 18684–18689. 10.1073/pnas.1110499108.

31. Zhang, H., Unal, H., Gati, C., Han, G.W., Liu, W., Zatsepin, N.A., James, D., Wang, D., Nelson, G., Weierstall, U., et al. (2015). Structure of the Angiotensin receptor revealed by serial femtosecond crystallography. Cell 161, 833–844. 10.1016/j.cell.2015.04.011.

32. Suomivuori, C.M., Latorraca, N.R., Wingler, L.M., Eismann, S., King, M.C., Kleinhenz, A.L.W., Skiba, M.A., Staus, D.P., Kruse, A.C., Lefkowitz, R.J., and Dror, R.O. (2020). Molecular mechanism of biased signaling in a prototypical G protein-coupled receptor. Science 367, 881–887. 10.1126/science.aaz0326.

33. Wingler, L.M., McMahon, C., Staus, D.P., Lefkowitz, R.J., and Kruse, A.C. (2019). Distinctive Activation Mechanism for Angiotensin Receptor Revealed by a Synthetic Nanobody. Cell 176, 479–490 e412. 10.1016/j.cell.2018.12.006.

34. Zhang, D., Liu, Y., Zaidi, S.A., Xu, L., Zhan, Y., Chen, A., Guo, J., Huang, X.P., Roth, B.L., Katritch, V., et al. (2023). Structural insights into angiotensin receptor signaling modulation by balanced and biased agonists. EMBO J. 42, e112940. 10.15252/embj.2022112940.

35. Zhang, H., Han, G.W., Batyuk, A., Ishchenko, A., White, K.L., Patel, N., Sadybekov, A., Zamlynny, B., Rudd, M.T., Hollenstein, K., et al. (2017). Structural basis for selectivity and diversity in angiotensin II receptors. Nature 544, 327–332. 10.1038/nature22035.

36. Yue, Y., Liu, L., Wu, L.J., Wu, Y., Wang, L., Li, F., Liu, J., Han, G.W., Chen, B., Lin, X., et al. (2022). Structural insight into apelin receptor-G protein stoichiometry. Nat. Struct. Mol. Biol. 29, 688–697. 10.1038/s41594-022-00797-5.

37. Chen, Q., and Tesmer, J.J.G. (2022). G protein-coupled receptor interactions with arrestins and GPCR kinases: The unresolved issue of signal bias. J. Biol. Chem. 298, 102279. 10.1016/j.jbc.2022.102279.

38. Staus, D.P., Wingler, L.M., Pichugin, D., Prosser, R.S., and Lefkowitz, R.J. (2019). Detergent- and phospholipid-based reconstitution systems have differential effects on constitutive activity of G-protein-coupled receptors. J. Biol. Chem. 294, 13218–13223. 10.1074/jbc.AC119.009848.

39. Lally, C.C., Bauer, B., Selent, J., and Sommer, M.E. (2017). C-edge loops of arrestin function as a membrane anchor. Nat. Commun. 8, 14258. 10.1038/ncomms14258.

40. Grimes, J., Koszegi, Z., Lanoiselee, Y., Miljus, T., O’Brien, S.L., Stepniewski, T.M., Medel-Lacruz, B., Baidya, M., Makarova, M., Mistry, R., et al. (2023). Plasma membrane preassociation drives beta-arrestin coupling to receptors and activation. Cell 186, 2238–2255 e2220. 10.1016/j.cell.2023.04.018.

41. Latorraca, N.R., Masureel, M., Hollingsworth, S.A., Heydenreich, F.M., Suomivuori, C.M., Brinton, C., Townshend, R.J.L., Bouvier, M., Kobilka, B.K., and Dror, R.O. (2020). How GPCR Phosphorylation Patterns Orchestrate Arrestin-Mediated Signaling. Cell 183, 1813–1825 e1818. 10.1016/j.cell.2020.11.014.

42. Maharana, J., Sarma, P., Yadav, M.K., Saha, S., Singh, V., Saha, S., Chami, M., Banerjee, R., and Shukla, A.K. (2023). Structural snapshots uncover a key phosphorylation motif in GPCRs driving beta-arrestin activation. Mol. Cell 83, 2091–2107 e2097. 10.1016/j.molcel.2023.04.025.

43. Cao, C., Barros-Alvarez, X., Zhang, S., Kim, K., Damgen, M.A., Panova, O., Suomivuori, C.M., Fay, J.F., Zhong, X., Krumm, B.E., et al. (2022). Signaling snapshots of a serotonin receptor activated by the prototypical psychedelic LSD. Neuron 110, 3154–3167 e3157. 10.1016/j.neuron.2022.08.006.

44. Lee, Y., Warne, T., Nehme, R., Pandey, S., Dwivedi-Agnihotri, H., Chaturvedi, M., Edwards, P.C., Garcia-Nafria, J., Leslie, A.G.W., Shukla, A.K., and Tate, C.G. (2020). Molecular basis of beta-arrestin coupling to formoterol-bound beta(1)-adrenoceptor. Nature 583, 862–866. 10.1038/s41586-020-2419-1.

45. Bous, J., Fouillen, A., Orcel, H., Trapani, S., Cong, X., Fontanel, S., Saint-Paul, J., Lai-Kee-Him, J., Urbach, S., Sibille, N., et al. (2022). Structure of the vasopressin hormone-V2 receptor-beta-arrestin1 ternary complex. Sci. Adv. 8, eabo7761. 10.1126/sciadv.abo7761.

46. Yin, W., Li, Z., Jin, M., Yin, Y.L., de Waal, P.W., Pal, K., Yin, Y., Gao, X., He, Y., Gao, J., et al. (2019). A complex structure of arrestin-2 bound to a G protein-coupled receptor. Cell Res. 29, 971–983. 10.1038/s41422-019-0256-2.

47. Kang, Y., Zhou, X.E., Gao, X., He, Y., Liu, W., Ishchenko, A., Barty, A., White, T.A., Yefanov, O., Han, G.W., et al. (2015). Crystal structure of rhodopsin bound to arrestin by femtosecond X-ray laser. Nature 523, 561–567. 10.1038/nature14656.

48. Aydin, Y., Bottke, T., Lam, J.H., Ernicke, S., Fortmann, A., Tretbar, M., Zarzycka, B., Gurevich, V.V., Katritch, V., and Coin, I. (2023). Structural details of a Class B GPCR-arrestin complex revealed by genetically encoded crosslinkers in living cells. Nat. Commun. 14, 1151. 10.1038/s41467-023-36797-2.

49. Bumbak, F., Bower, J.B., Zemmer, S.C., Inoue, A., Pons, M., Paniagua, J.C., Yan, F., Ford, J., Wu, H., Robson, S.A., et al. (2023). Stabilization of pre-existing neurotensin receptor conformational states by beta-arrestin-1 and the biased allosteric modulator ML314. Nat. Commun. 14, 3328. 10.1038/s41467-023-38894-8.

50. Xu, J., Wang, Q., Hubner, H., Hu, Y., Niu, X., Wang, H., Maeda, S., Inoue, A., Tao, Y., Gmeiner, P., et al. (2023). Structural and dynamic insights into supra-physiological activation and allosteric modulation of a muscarinic acetylcholine receptor. Nat. Commun. 14, 376. 10.1038/s41467-022-35726-z.

51. Seyedabadi, M., Gharghabi, M., Gurevich, E.V., and Gurevich, V.V. (2022). Structural basis of GPCR coupling to distinct signal transducers: implications for biased signaling. Trends Biochem. Sci. 47, 570–581. 10.1016/j.tibs.2022.03.009.

52. Zhao, J., Elgeti, M., O’Brien, E.S., Sar, C.P., Ei Daibani, A., Heng, J., Sun, X., White, E., Che, T., Hubbell, W.L., et al. (2024). Ligand efficacy modulates conformational dynamics of the micro-opioid receptor. Nature 629, 474–480. 10.1038/s41586-024-07295-2.

53. Staus, D.P., Wingler, L.M., Choi, M., Pani, B., Manglik, A., Kruse, A.C., and Lefkowitz, R.J. (2018). Sortase ligation enables homogeneous GPCR phosphorylation to reveal diversity in beta-arrestin coupling. Proc. Natl. Acad. Sci. U.S.A 115, 3834–3839. 10.1073/pnas.1722336115.

54. Kawakami, K., Yanagawa, M., Hiratsuka, S., Yoshida, M., Ono, Y., Hiroshima, M., Ueda, M., Aoki, J., Sako, Y., and Inoue, A. (2022). Heterotrimeric Gq proteins act as a switch for GRK5/6 selectivity underlying beta-arrestin transducer bias. Nat. Commun. 13, 487. 10.1038/s41467-022-28056-7.

55. Drube, J., Haider, R.S., Matthees, E.S.F., Reichel, M., Zeiner, J., Fritzwanker, S., Ziegler, C., Barz, S., Klement, L., Filor, J., et al. (2022). GPCR kinase knockout cells reveal the impact of individual GRKs on arrestin binding and GPCR regulation. Nat. Commun. 13, 540. 10.1038/s41467-022-28152-8.

56. Wei, H., Ahn, S., Barnes, W.G., and Lefkowitz, R.J. (2004). Stable interaction between beta-arrestin 2 and angiotensin type 1A receptor is required for beta-arrestin 2-mediated activation of extracellular signal-regulated kinases 1 and 2. J. Biol. Chem. 279, 48255–48261. 10.1074/jbc.M406205200.

57. Zimmerman, B., Beautrait, A., Aguila, B., Charles, R., Escher, E., Claing, A., Bouvier, M., and Laporte, S.A. (2012). Differential beta-arrestin-dependent conformational signaling and cellular responses revealed by angiotensin analogs. Sci. Signal. 5, ra33. 10.1126/scisignal.2002522.

58. Santos, G.A., Duarte, D.A., Parreiras, E.S.L.T., Teixeira, F.R., Silva-Rocha, R., Oliveira, E.B., Bouvier, M., and Costa-Neto, C.M. (2015). Comparative analyses of downstream signal transduction targets modulated after activation of the AT1 receptor by two beta-arrestin-biased agonists. Front. Pharmacol. 6, 131. 10.3389/fphar.2015.00131.

59. Liu, Q., He, Q.T., Lyu, X., Yang, F., Zhu, Z.L., Xiao, P., Yang, Z., Zhang, F., Yang, Z.Y., Wang, X.Y., et al. (2020). DeSiphering receptor core-induced and ligand-dependent conformational changes in arrestin via genetic encoded trimethylsilyl (1)H-NMR probe. Nat. Commun. 11, 4857. 10.1038/s41467-020-18433-5.

60. Petrovic, I., Tatli, M., Desai, S., Grahl, A., Ni, D., Stahlberg, H., Spang, A., Grzesiek, S., and Abiko, L.A. (2025). Arrestin recognizes GPCRs independently of the receptor state. Proc. Natl. Acad. Sci. U.S.A 122, e2501487122. 10.1073/pnas.2501487122.

61. Xu, J., Hu, Y., Kaindl, J., Risel, P., Hubner, H., Maeda, S., Niu, X., Li, H., Gmeiner, P., Jin, C., and Kobilka, B.K. (2019). Conformational Complexity and Dynamics in a Muscarinic Receptor Revealed by NMR Spectroscopy. Mol. Cell 75, 53–65 e57. 10.1016/j.molcel.2019.04.028.

62. Staus, D.P., Strachan, R.T., Manglik, A., Pani, B., Kahsai, A.W., Kim, T.H., Wingler, L.M., Ahn, S., Chatterjee, A., Masoudi, A., et al. (2016). Allosteric nanobodies reveal the dynamic range and diverse mechanisms of G-protein-coupled receptor activation. Nature 535, 448–452. 10.1038/nature18636.

63. Pannier, M., Veit, S., Godt, A., Jeschke, G., and Spiess, H.W. (2000). Dead-time free measurement of dipole-dipole interactions between electron spins. J Magn Reson 142, 331–340. 10.1006/jmre.1999.1944.

64. Altenbach, C. (2021). LongDistances—a program to analyze DEER data. EPR Newsletter 31, 12–13.

65. Jumper, J., and Hassabis, D. (2022). Protein structure predictions to atomic accuracy with AlphaFold. Nat. Methods 19, 11–12. 10.1038/s41592-021-01362-6.

66. Lomize, M.A., Pogozheva, I.D., Joo, H., Mosberg, H.I., and Lomize, A.L. (2012). OPM database and PPM web server: resources for positioning of proteins in membranes. Nucleic Acids Res. 40, D370–376. 10.1093/nar/gkr703.

67. Jo, S., Kim, T., Iyer, V.G., and Im, W. (2008). CHARMM-GUI: a web-based graphical user interface for CHARMM. J. Comput. Chem. 29, 1859–1865. 10.1002/jcc.20945.

68. Van Der Spoel, D., Lindahl, E., Hess, B., Groenhof, G., Mark, A.E., and Berendsen, H.J. (2005). GROMACS: fast, flexible, and free. J. Comput. Chem. 26, 1701–1718. 10.1002/jcc.20291.

69. Berendsen, H.J.C., Postma, J.P.M., Vangunsteren, W.F., Dinola, A., and Haak, J.R. (1984). Molecular-Dynamics with Coupling to an External Bath. J. Chem. Phys. 81, 3684–3690. 10.1063/1.448118.

70. Parrinello, M., and Rahman, A. (1981). Polymorphic Transitions in Single-Crystals - a New Molecular-Dynamics Method. J. Appl. Phys. 52, 7182–7190. 10.1063/1.328693.

71. Bussi, G., Donadio, D., and Parrinello, M. (2007). Canonical sampling through velocity rescaling. J. Chem. Phys. 126, 014101. 10.1063/1.2408420.

72. Darden, T., York, D., and Pedersen, L. (1993). Particle Mesh Ewald - an N.Log(N) Method for Ewald Sums in Large Systems. J. Chem. Phys. 98, 10089–10092. 10.1063/1.464397.

73. Essmann, U., Perera, L., Berkowitz, M.L., Darden, T., Lee, H., and Pedersen, L.G. (1995). A Smooth Particle Mesh Ewald Method. J. Chem. Phys. 103, 8577–8593. 10.1063/1.470117.

74. Hess, B. (2008). P-LINCS: A Parallel Linear Constraint Solver for Molecular Simulation. J. Chem. Theory Comput. 4, 116–122. 10.1021/ct700200b.

75. Weis, W.I., and Kobilka, B.K. (2018). The Molecular Basis of G Protein-Coupled Receptor Activation. Annu. Rev. Biochem. 87, 897–919. 10.1146/annurev-biochem-060614-033910.

76. Dalton, J.A., Lans, I., and Giraldo, J. (2015). Quantifying conformational changes in GPCRs: glimpse of a common functional mechanism. BMC Bioinformatics 16, 124. 10.1186/s12859-015-0567-3.

77. Gowers, R.J., Linke, M., Barnoud, J., Reddy, T.J.E., Melo, M.N., Seyler, S.L., Domanski, J., Dotson, D.L., Buchoux, S., Kenney, I.M., and Beckstein, O. (2016). MDAnalysis: A Python Package for the Rapid Analysis of Molecular Dynamics Simulations. In S. Benthall, and S. Rostrup, eds. Proceedings of the 15th Python in Science Conference. SciPy.

78. Michaud-Agrawal, N., Denning, E.J., Woolf, T.B., and Beckstein, O. (2011). MDAnalysis: a toolkit for the analysis of molecular dynamics simulations. J. Comput. Chem. 32, 2319–2327. 10.1002/jcc.21787.

79. Tessmer, M.H., and Stoll, S. (2023). chiLife: An open-source Python package for in silico spin labeling and integrative protein modeling. PLoS Comput Biol 19, e1010834. 10.1371/journal.pcbi.1010834.

80. Elgeti, M., Rose, A.S., Bartl, F.J., Hildebrand, P.W., Hofmann, K.P., and Heck, M. (2013). Precision vs flexibility in GPCR signaling. J. Am. Chem. Soc. 135, 12305–12312. 10.1021/ja405133k.

