## Supplemental information for "Angiotensin receptor conformations stabilized by biased ligands differentially modulate β-arrestin interactions"

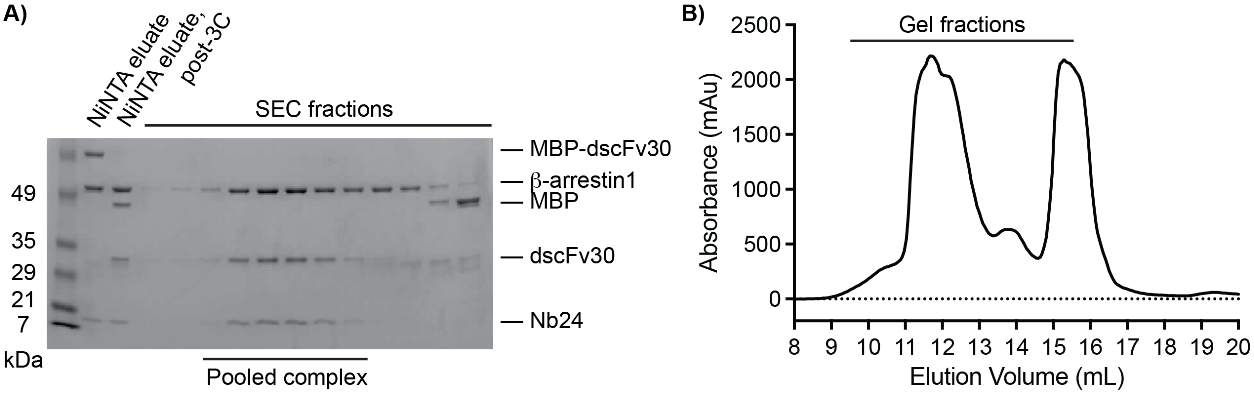


**Figure S1.** Purification of βarr1*. **A)** Gel electrophoresis analysis (Coomassie staining) of βarr1* complex following NiNTA purification, 3C cleavage, and size exclusion chromatography. SEC fractions pooled for subsequent experiments are indicated. **B)** SEC purification of βarr1*. Fractions analyzed by gel electrophoresis are indicated.

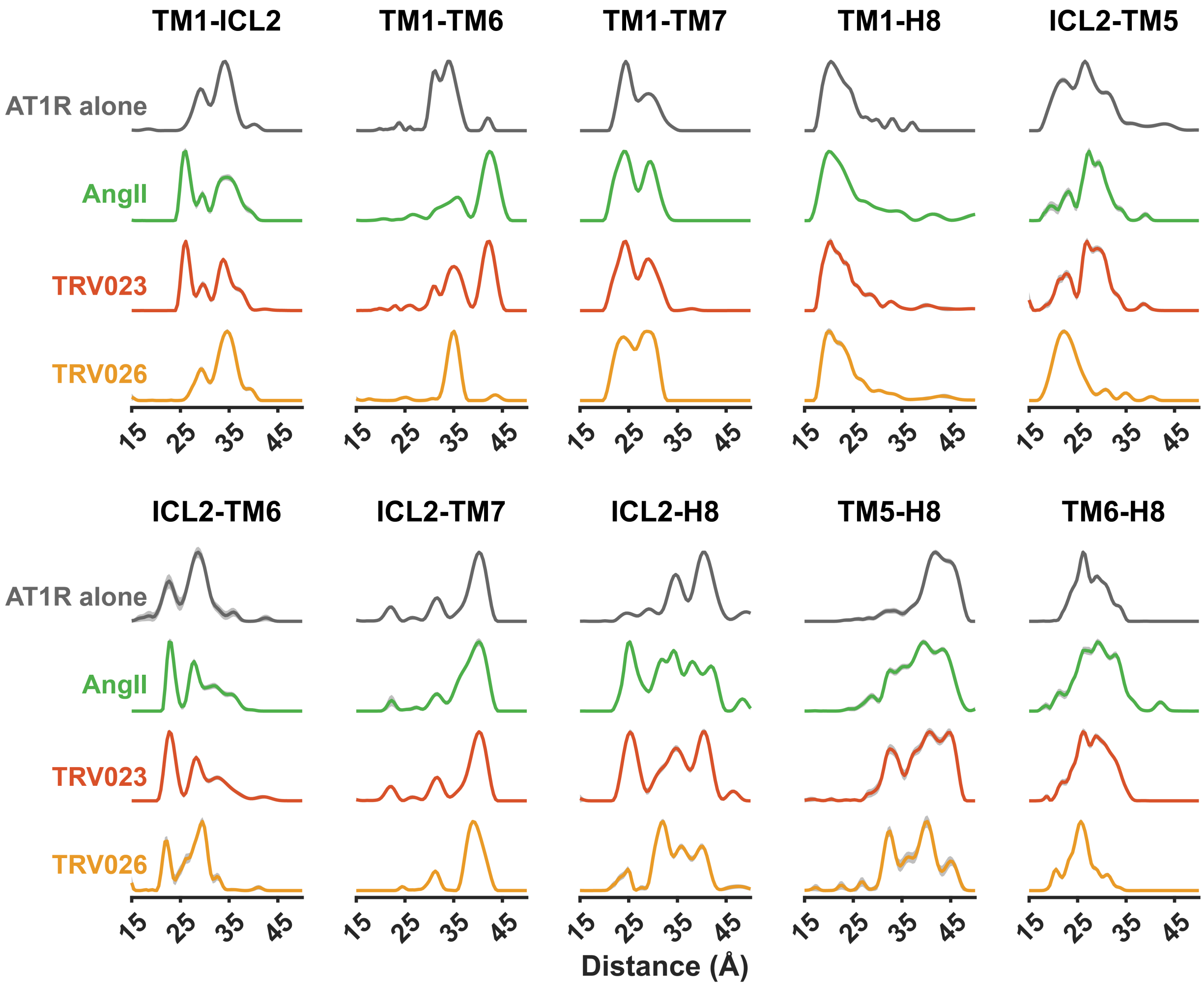


**Figure S2.** Probability distance distributions indicating the distinct effects of AngII (green), TRV023 (red-orange) and TRV026 (light orange) binding compared to unliganded AT1R (gray). Shaded areas indicate 95% confidence intervals for the model-free analysis.

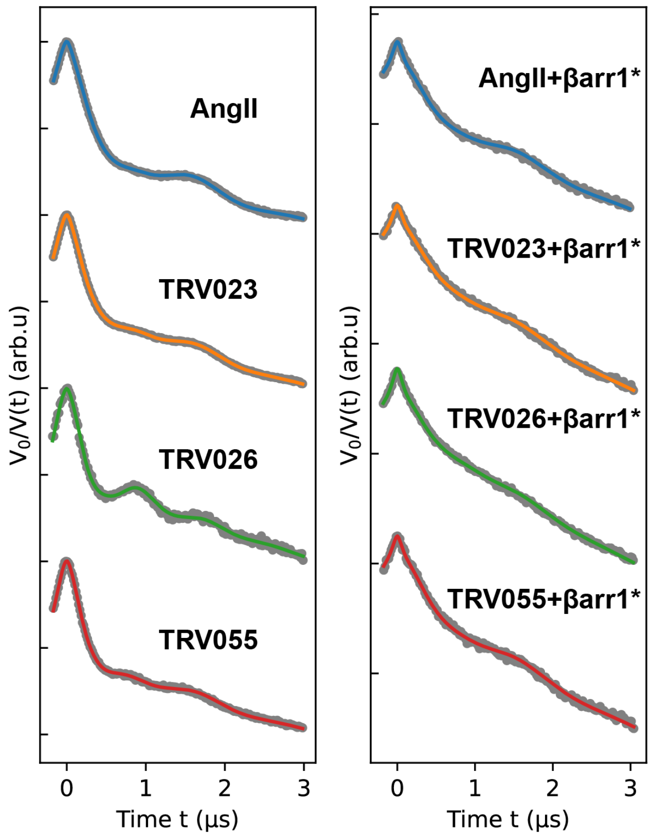


**Figure S3.** Raw DEER data (gray) and model-based dipolar analyses (colored) for the TM1-TM6 pair under the indicated ligand conditions.

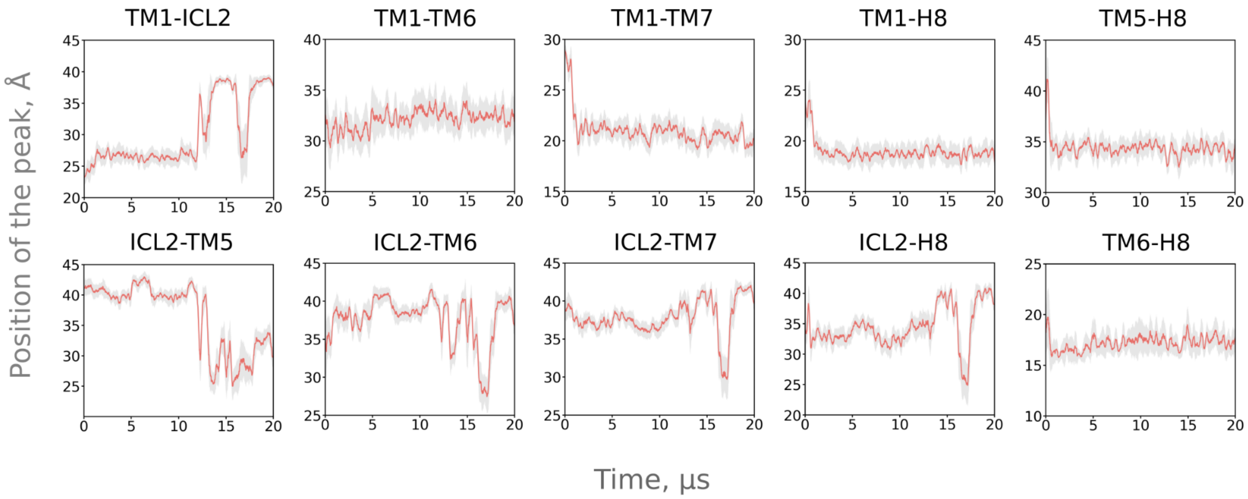


**Figure S4**. Position of the peak of the simulated distance distributions over time. The red line represents the sliding mean, while the semi-transparent gray area indicates the standard deviation.

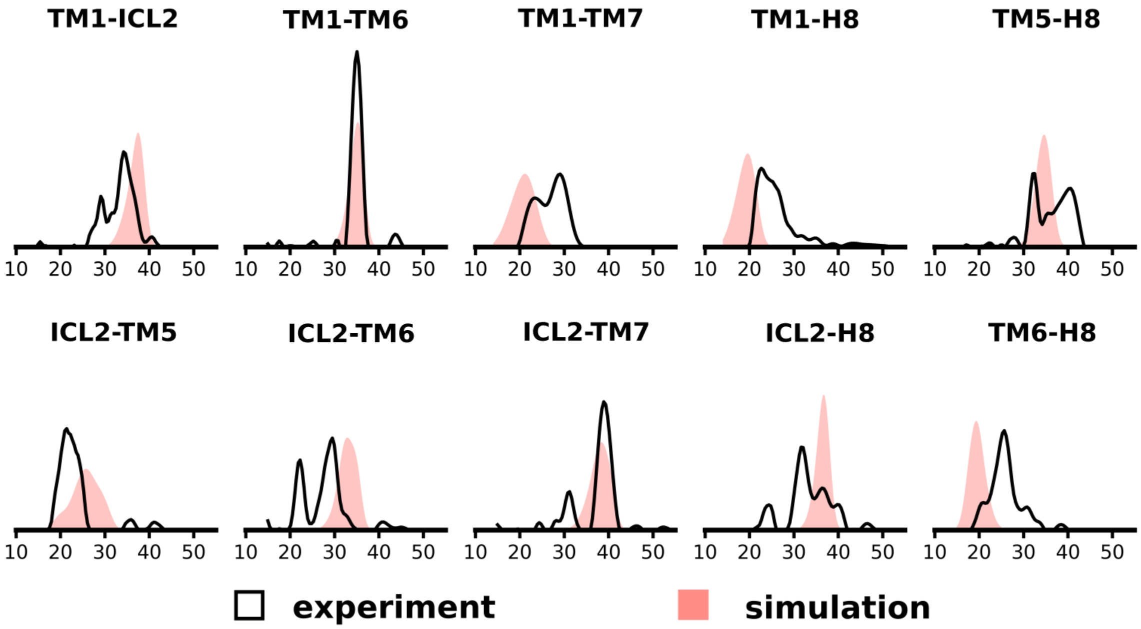


**Figure S5**. Direct comparison of distance distributions from DEER and unbiased MD simulations.

**Table S1**

| | | **Log $IC_{50}$ (95% CI)** |
|---|---|---|
| **AngII** | **-βarr1*** | -7.14 (-7.22 – -7.07) |
| | **+βarr1*** | -7.61 (-7.78 – -7.44) |
| **Losartan** | **-βarr1*** | -6.55 (-6.69 – -6.40) |
| | **+βarr1*** | -6.68 (-6.79 – -6.58) |

Log $IC_{50}$ values for competition radioligand binding data shown in Fig. 1B,C.

**Table S2**

| Ligand | Log $EC_{50}$ (95% CI) | $E_{max}$ (95% CI) |
|---|---|---|
| **AngII** | -8.15 (-8.25 – -8.05) | 100 (96 – 103) |
| **TRV023** | -7.45 (-7.52 – -7.37) | 53 (52 – 55) |
| **TRV027** | -7.62 (-7.70 – -7.54) | 48 (47 – 49) |
| **TRV026** | -7.67 (-7.76 – -7.58) | 60 (58 – 62) |
| **Candesartan** | N.R. | N.R. |
| **Losartan** | N.R. | N.R. |
| **Olmesartan** | N.R. | N.R. |

Log $EC_{50}$ and $E_{max}$ values for ligand-induced β-arrestin1 plasma membrane recruitment, including the data shown in Fig. 3A. N.R., no response.

**Table S3**

| | | 23 Å | 32 Å | 35 Å | 42 Å |
|---|---|---|---|---|---|
| **Apo** | **-βarr1*** | 0.083 (0.072 – 0.094) | 0.534 (0.508 – 0.559) | 0.268 (0.244 – 0.293) | 0.115 (0.105 – 0.125) |
| | **+βarr1*** | N.D. | N.D. | N.D. | N.D. |
| **AngII** | **-βarr1*** | 0.061 (0.054 – 0.069) | 0.096 (0.082 – 0.110) | 0.173 (0.160 – 0.186) | 0.672 (0.665 – 0.678) |
| | **+βarr1*** | 0.261 (0.223 – 0.299) | 0.009 (0.00e+00 – 0.079) | 0.00e+00 (0.00e+00 – 0.056) | 0.730 (0.704 – 0.755) |
| **TRV023** | **-βarr1*** | 0.058 (0.052 – 0.063) | 0.161 (0.149 – 0.174) | 0.288 (0.277 – 0.300) | 0.494 (0.488 – 0.499) |
| | **+βarr1*** | 0.282 (0.205 – 0.358) | 0.00e+00 (0.00e+00 – 0.119) | 0.00e+00 (0.00e+00 – 0.075) | 0.721 (0.672 – 0.769) |
| **TRV027** | **-βarr1*** | 0.049 (0.041 – 0.057) | 0.154 (0.138 – 0.171) | 0.375 (0.360 – 0.390) | 0.423 (0.416 – 0.429) |
| | **+βarr1*** | N.D. | N.D. | N.D. | N.D. |
| **TRV026** | **-βarr1*** | 0.031 (0.003 – 0.059) | 0.097 (0.043 – 0.151) | 0.832 (0.783 – 0.881) | 0.039 (0.017 – 0.062) |
| | **+βarr1*** | 0.335 (0.286 – 0.384) | 0.00e+00 (0.00e+00 – 0.133) | 0.133 (0.026 – 0.240) | 0.537 (0.508 – 0.566) |
| **TRV055** | **-βarr1*** | 0.040 (0.030 – 0.050) | 0.321 (0.297 – 0.345) | 0.253 (0.230 – 0.275) | 0.388 (0.378 – 0.397) |
| | **+βarr1*** | 0.234 (0.186 – 0.281) | 0.00e+00 (0.00e+00 – 0.120) | 0.00e+00 (0.00e+00 – 0.133) | 0.758 (0.723 – 0.792) |

Gaussian populations (median ± 95% confidence interval) centered around the indicated TM1-TM6 distances for data shown in Fig. 3C.

**Table S4**

| # | PDB ID | Arrestin | GPCR |
|---|---|---|---|
| 1 | 6UP7 | β-arrestin 1 | NT1R |
| 2 | 8WRZ | β-arrestin 1 | CB1R |
| 3 | 8ZYT | β-arrestin 1 | NT1R |
| 4 | 8ZYY | β-arrestin 1 | NT1R |
| 5 | 7R0C | β-arrestin 1 | V2R |
| 6 | 6U1N | β-arrestin 1 | M2R/V2R chimera |
| 7 | 6PWC | β-arrestin 1 | NT1R |
| 8 | 6UP7 | β-arrestin 1 | NT1R |
| 9 | 6TKO | β-arrestin 1 | β1AR |
| 10 | 7SRS | β-arrestin 1 | 5-HT2BR |
| 11 | 8WU1 | β-arrestin 1 | CB1R |
| 12 | 9M0D | β-arrestin 1 | NT1R |
| 13 | 8ZYU | β-arrestin 1 | NT1R |
| 14 | 5DGY | visual arrestin | rhodopsin |
| 15 | 5W0P | visual arrestin | rhodopsin |
| 16 | 4ZWJ | visual arrestin | rhodopsin |

GPCR-arrestin complexes used for the alignment in Fig. 5C.
